# Sin3a Associated Protein 130kDa, sap130, plays an evolutionary conserved role in zebrafish heart development

**DOI:** 10.1101/2023.03.30.534737

**Authors:** Ricardo A. DeMoya, Rachel E. Forman-Rubinsky, Deon Fontaine, Joseph Shin, Simon C. Watkins, Cecilia Lo, Michael Tsang

## Abstract

Hypoplastic left heart syndrome (HLHS) is a congenital heart disease where the left ventricle is reduced in size. A forward genetic screen in mice identified SIN3A associated protein 130kDa (*Sap130*), a protein in the chromatin modifying SIN3A/HDAC1 complex, as a gene contributing to the digenic etiology of HLHS. Here, we report the role of zebrafish *sap130* genes in heart development. Loss of *sap130a,* one of two *Sap130* orthologs, resulted in smaller ventricle size, a phenotype reminiscent to the hypoplastic left ventricle in mice. While cardiac progenitors were normal during somitogenesis, diminution of the ventricle size suggest the Second Heart Field (SHF) was the source of the defect. To explore the role of *sap130a* in gene regulation, transcriptome profiling was performed after the heart tube formation to identify candidate pathways and genes responsible for the small ventricle phenotype. Genes involved in cardiac differentiation and cell communication were dysregulated in *sap130a*, but not in *sap130b* mutants. Confocal light sheet analysis measured deficits in cardiac output in *MZsap130a* supporting the notion that cardiomyocyte maturation was disrupted. Lineage tracing experiments revealed a significant reduction of SHF cells in the ventricle that resulted in increased outflow tract size. These data suggest that *sap130a* is involved in cardiogenesis via regulating the accretion of SHF cells to the growing ventricle and in their subsequent maturation for cardiac function. Further, genetic studies revealed an interaction between *hdac1* and *sap130a*, in the incidence of small ventricles. These studies highlight the conserved role of Sap130a and Hdac1 in zebrafish cardiogenesis.

## Introduction

Congenital heart diseases (CHDs) affect approximately 1% of live births per year and causes have been attributed to environmental and genetic factors (1, 2, 3, 4). Hypoplastic left heart syndrome (HLHS) is a severe CHD characterized by a reduced volume in the left ventricle and aortic and valve malformations (5, 6). The genetic etiology of HLHS is complex and genetically heterogenous. Mouse models of HLHS were recovered from a large-scale mutagenesis screen (7), and among 8 lines, the *Ohia* mutant line was identified to have a digenic etiology for HLHS. This is comprised of mutations in SIN3A associated protein 130kDa (SAP130) and protocadherin 9 (PCDHA9) that together causes HLHS comprising hypoplasia of all left-sided heart structures including the ventricle, aorta/aortic valve, and mitral valve. In pigs a CRISPR generated SAP130 allele caused embryonic lethality and tricuspid dysplasia and atresia, indicating SAP130 involvement in cardiac development in higher vertebrates (8). In zebrafish a maternal zygotic *sap130a* (*MZsap130a)* mutant produced a diminutive ventricle by 72 hours post fertilization (hpf), confirming that SAP130 retains a conserved function among vertebrates during heart development (7, 8).

SAP130 was identified as an interacting protein in the SIN3A complex, binding both SIN3A and Histone Deacetylase 1 (HDAC1), thought to stabilize the complex. It was theorized that the SAP130 C-terminus functioned as a transcriptional repressor in association with the SIN3A complex, while the N-terminus paradoxically could function as an activator (9). A knock-out allele of SAP130 in mice is lethal, similar to global HDAC1 and SIN3A knockouts (7, 8, 10, 11). SIN3A and HDACs epigenetically regulate gene transcription through histone and non-histone deacetylation events and are classically associated with gene repression. However, some studies have shown this complex to be a transcriptional activator in other contexts (10, 12, 13). HDACs have been reported to regulate many aspects of development, including cardiac development in zebrafish, mouse, and chick models, evidenced by treatment with a pan HDAC small molecule inhibitor, Trichostatin A (14, 15, 16, 17). Zebrafish studies have revealed that *hdac1* is involved in SHF heart development and adult cardiac regeneration (18, 19). In zebrafish, *hdac1* mutants have less cardiomyocytes (CMs) in the ventricle while inhibition of *hdac1* (and other class I HDACs) reveal reduced proliferation during regenerative events (18, 19, 20, 21). Zebrafish *hdac1* mutants are embryonic lethal, similar to the mouse models, but *MZsap130a* mutants are viable as adults suggesting that *hdac1* and *sap130a* may have distinct functions in zebrafish cardiogenesis (18, 22). In addition, members of the SIN3 complex have been shown to be involved in myotube differentiation through regulation of sarcomere gene expression. A SIN3A/B knock-down study in cultured myotubes decreased sarcomere genes actin (*Acta1)*, Titin (*Ttn)*, Tropinin-C1 (*Tnnc1)* and Tropomyosin4 (*Tpm4)*, suggesting the SIN3A complex playing a role in CM differentiation (23, 24, 25). Zebrafish *sin3b* mutants are viable but showed only skeletal defects and *sin3aa* or *sin3ab* knock-down studies revealed involvement in hematopoiesis (26, 27).

In addition to the Sin3a/Hdac1 complex, other mutations in genes that function as chromatin modifiers such as Brahma-related gene 1 (*brg1)* and SET and MYND domain-containing lysine methyltransferase 4 (*smyd4),* also result in reduced ventricle size in zebrafish and mouse, suggesting there is a common requirement of gene regulation for specifying heart organ size (28). Zebrafish *brg1* mutants have reduce CM proliferation leading to a smaller ventricle after 28hpf. The *brg1* mutants reveal changes in a working myocardium marker *nppa*, similar to mouse Brg1 mutants (29). *Smyd4*, another epigenetic regulator, has been shown to be involved in zebrafish heart development. RNA sequencing (RNA-seq) analysis of *smyd4* zebrafish mutants revealed dysregulation of cardiac muscle contraction genes and metabolism. Moreover, cell culture studies revealed human SMYD4 and HDAC1 interact, further supporting a central requirement for hdac1 in zebrafish cardiogenesis (30). Taken together these suggest a potential epigenetic role for *sap130a* during cardiogenesis.

Zebrafish are an exceptional model to study cardiac development. Greater than seventy percent of human protein coding genes having at least one zebrafish orthologue and eighty percent of those genes are disease related (31). Several studies have highlighted the similarities between zebrafish and mammalian cardiogenesis (32, 33, 34, 35, 36, 37). Zebrafish cardiac progenitors can be first delineated in the late blastula stage (5hpf, shield stage) and are located at the lateral marginal zone intermingled with other mesoderm lineages (38, 39). By 15hpf (12 somite stage) the cardiac progenitor cells have migrated to the lateral plate mesoderm. Subsequently these bilateral cardiac populations coalesce to fuse into a heart tube and begin to differentiate into cardiomyocytes. At 18hpf to 20hpf (18s-22s), the progenitors surround the endocardial cells and form a cardiac disk with a lumen in the center (40). From 20hpf to 36hpf (22s to prim-22) the disk will elongate in an anterior direction to form the heart tube, which makes up the first heart field (FHF) and undergoes jogging and ballooning processes. By 48hpf (long-pec) the second heart field (SHF) is added and the tube moves left and antero-ventrally until the ventricle is at ventral midline and atria in a right-ventral position (41, 42, 43). The early SHF cells remain bilateral populations of CMs at first and then follow the heart tube as it positions itself, continuously adding cells to the ventricle and finally the OFT (38, 44, 45, 46, 47, 48, 49, 50, 51). All these steps in cardiac development are epigenetically regulated and are dependent on many early developmental pathways, including Retinoic Acid, fibroblast growth factors, Wnt, and bone morphogenic proteins (38, 39, 52, 53, 54, 55, 56). As these major pathways are also critical for early developmental patterning, their distinct function in cardiogenesis has been difficult to ascertain in the zebrafish embryo.

Here we investigate the role of *sap130* genes in zebrafish by studying mutations in both *sap130a* and *sap130b*. Transcriptome profiling of 36hpf *MZsap130a* mutants revealed over 5000 genes to be differentially expressed, including genes involved in cardiac sarcomere assembly. In genetic studies, an increase in embryos with small ventricles (SVs) were noted in *MZsap130a* embryos that were also heterozygous for *hdac1*. Furthermore, *MZsin3ab* mutant embryos also exhibit an SV phenotype. Collectively, these studies suggest a role for *sin3ab*/*hdac1*/*sap130a* in SHF cardiomyocyte maturation and communication in zebrafish heart development.

## Materials and Methods

### Zebrafish Husbandry

All zebrafish experiments and protocols were performed according to protocols approved by the Institutional Animal Care and Use Committee (IACUC) at the University of Pittsburgh in agreement with NIH guidelines. Wild-type *AB\**, *Tg(myl7:GFP)^twu34^* (57), *Tg(nkx2.5:kaede)^fb9^*(45), *sap130a^pt32a^*(7), *hdac1^b382^*(22)

Adult tail fin clips or whole embryos for genotyping assays was performed as previously described (58). Restriction fragment length polymorphism (RFLP) genotyping for *sap130a^pt32a^*, *sap130b^pt35b^*, *sin3ab^pt36a^* and *hdac1^b382^* used the primers and enzymes listed in **Supplementary Table S1**.

### CRISPR/Cas9 mutant allele generation

The CRISPR/Cas9 protocol (59) was used to establish mutant lines. This protocol used Sp6 in vitro transcribed sgRNAs targeting the sequence ccgTGGGAGGGAAAACAATGCTG for *sap130b* and cctGCTCCTCTTCAGCCATACAG for *sin3ab*, where lower case letters represent the protospacer motif sequence. sgRNA was incubated at room temperature with Cas9 protein (NEB, Cat# M0646T). *AB\** embryos were injected at the one-cell stage with the sgRNA and Cas9 cocktail in a 1nL volume at 25pg sgRNA/nL. RFLP was performed to determine protected mutated bands present 24hrs after injection to determine gRNA efficiency and injected embryos were raised to adults outcrossed to *AB\**. DNA mutations in *sap130b* and *sin3ab* were verified by PCR TOPO-TA cloning (ThermoFisher, #K4575J10) from adult heterozygous animals and Sanger sequenced. gRNA sequence information **Supplementary Table S2** and **Supplementary Table S3**.

### Imaging

A Leica M205 FA stereomicroscope was used to take images of the hearts from *Tg(myl7:EGFP)* WT and mutant embryos at 36 and 48hpf. For imaging the *Tg(myl7:memGFP)* OFT, a Nikon A1 inverted confocal microscope was used at 72hpf. *Tg(myl7:memGFP)* embryos were anesthetized in 7x MS-222/10mM BDM (2,3-butanedione monoxime) and mounted in low melting agarose on MaTek glass bottom petri dish (MaTek, Part No: P35G-1.5-14-C) and imaged at a 40x water immersion.

### ConSurf and R generated phylogenetic trees and protein diagram

ConSurf (https://consurf.tau.ac.il/consurf_index.php) was used to align multiple Sap130 protein sequences across many species (60). The *sap130a* amino acid sequence from zebrafish was input to ConSurf and the output was collected and plotted in R, with ggtree, ggplot2 and phytools (61, 62, 63, 64). A multiple sequence alignment (MSA) was performed on Sap130 protein sequences from UniProt and distance calculations to plot simple phylogeny trees using R CRAN packages seqinr, msa, Biostrings, ggtree, ggplot2 (65, 66, 67). For plotting the protein sequences and conserved domains reported by UniProt, the R packages ggplot and drawProteins were used (68).

### *in situ* probe synthesis, whole mount *in situ* hybridization

RNA probe generation and whole mount in situ hybridization for *nkx2.5*, *myh7* and *myh6* was performed as previously described with DIG RNA labeling kit (Millipore Sigma cat# 11175025910) (69)

### RNAseq sample preparation and data analysis

Total RNA was extracted from whole embryos or isolated hearts (36hpf and 48hpf, respectively) using Trizol (Invitrogen) and was purified with the RNeasy Micro Kit (Qiagen#74004). A minimum 50 embryos or 180 hearts were pooled together for each condition. The RNA-seq used was 0.5-1μg RNA for each condition and was sent to the Genomics Research Core at the University of Pittsburgh. The raw sequence reads were processed and mapped to the Zebrafish Reference Genome GRCz11 using CLC Genomics Workbench 20. A count matrix was exported and the bioinformatic analysis was carried out in R (70) using the edgeR package for 36hpf whole embryo and 48hpf heart tissue data. Results for DEGs in **Supplementary Tables, S4, S5, S6, S7, S8** (71). After determining differentially expressed genes they were entered into DAVID (https://david.ncifcrf.gov/summary.jsp) for functional annotation clustering. Results for DAVID clustering in **Supplementary Tables, S4, S5, S8** (72).

### Lineage Tracing

Lineage tracing of cardiac progenitors at 24hpf was performed on *Tg(nkx2.5: kaede)* and *Tg(nkx2.5:kaede);sap130a^pt32a/pt32a^* embryos was described by Guner-Ataman et al (45). Using the Zeiss Imager M2 confocal microscope at 40x, the ROI (Region of Interest) was selected to photoconvert the peristaltic heart tube at 24hpf. Embryos were mounted in low melting temperature agarose droplets on 35mm dishes. The embryos were then freed from the agarose and raised in darkness until 48hpf, when the looped heart was imaged at 40x.

### Cardiac functional analysis

To measure cardiac function in embryonic zebrafish, we used a custom-built light sheet microscope which followed a design based on the openSPIM platform (73) (74). This ‘T’ design illuminates the sample bilaterally and uses a four-channel laser launch for maximum versatility. *Tg(myl7:EGFP)* and *Tg(myl7:EGFP);sap130a^m/m^* embryos at 48hpf embryos were placed into E3 and Tricaine (307 nmol concentration) to anesthetize them before mounting for imaging. Low melting point agarose was heated and cooled to 42°C. 100µL agarose placed onto a dish and after 45 seconds of cooling, 48hpf embryo was added to the agarose and drawn into a custom cut 1 ml straight-barreled syringe. The agarose is allowed to solidify, and the syringe is placed into a sample manipulator capable of 3D movement + rotation (Picard Technologies, Inc.). The agarose-embedded embryos were extruded from the syringe and positioned in a lateral view, with anterior to the left and posterior to the right, before recording 100 frames at 50-75 frames per second using a Prime 95B sCMOS camera (Photometrics, Inc.). Fiji ImageJ software was used to identify end-diastole and end-systole frames to calculate ventricle area, length (distance between ventricular apex and out-flow tract opening), and diameter for each embryo (distance between the walls of the chamber, taken from the middle of length measurement). These data were used to estimate chamber volumes and calculate end-diastole and systole volumes, ejection fraction (%), fractional shortening (µm), Total stroke volume, cardiac output, and heart rate as an average of all cycles captured for each fish. The volumes calculated are under the assumption of a prolate sphere shape (p/6). The equations used are as follows (75);

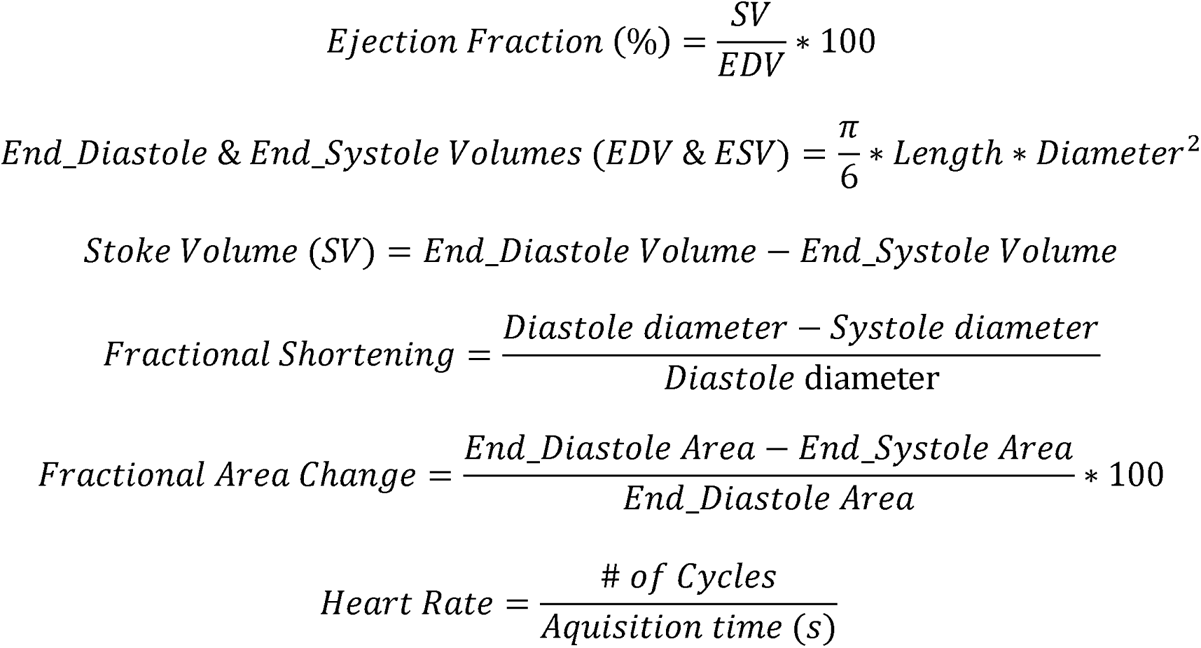

These were implemented using R scripting and RStudio to automate the calculations, and then data were plotted using Graphpad PRISM9. Each data point represents an average of 3 or more cycles per fish (75).

### Adult heart measurements

At 48hpf *MZsap130a* mutant embryos were scored for ventricle size and raised in separate tanks. *MZsap130a* mutants and aged matched *AB\** controls were measured for length and weight before hearts were extracted for DIC imaging at 4-6mpf. Fiji-ImageJ was used to measure the ventricle surface area and BA surface area. These data were plotted using Graphpad Prism9.

### Statistics

For analysis of RNA-seq data we used the edgeR package, utilizing a quasi-likelihood negative binomial generalized log-linear model to our count data comparing AB* control to *MZsap130a* or *MZsap130b* mutant embryos at 36hpf. For heart tissue RNA-seq, edgeR’s likelihood ratio test was used to interpret up or down regulation of genes. For the all other statistical analysis, significance was calculated using two-tailed, unpaired Student’s t-test, one-way ANOVA or Fisher’s exact text using GraphPad Prism version 9.3.

## Results

### *sap130b* is not required for heart development

Zebrafish were part of the teleost-specific genome duplication event 350 million years ago (76), resulting in two sap130 genes, *sap130a* and *sap130b*. Defining the SAP130 protein domains based on homology with other model organisms will provide insight into the potential conserved functional domains. In mammals, both SIN3A and HDAC1 proteins were shown to interact with SAP130 at the C-terminus between amino acids 836-1047, suggesting that SAP130 may act as a stabilizing scaffold between these proteins (9). Determining protein sequence similarities can predict functional structures across species and offer insight into the potential for functional redundancy between Sap130a and Sap130b. ConSurf was used for a multispecies comparison of 145 SAP130 protein sequences to determine their similarity and conserved domains (60). In general, Sap130a and Sap130b are dissimilar, but they both contained conserved N- and C-terminus domains represented by repetitive predicted structural and functional residues (**Figure S1**). When comparing a smaller set of protein sequences among other teleost, Sap130a and Sap130b remain different suggesting these dissimilarities are consistent with other species (**Figure 1A**). However, when compared to a broader group, these two proteins are most similar to one another (**Figure 1B**). This suggests that Sap130a and Sap130b share similar domains and can potentially compensate for one another in zebrafish. *MZsap130a* mutants develop SVs in 36% of the population by 72hpf (7). The incomplete penetrance of the SV phenotype was hypothesized to be the result of *sap130b* compensation for the loss of *sap130a*. To address this, we generated a mutation in *sap130b* using CRISPR/Cas9 technology. This produced an allele (7bp del,1bp sub (G>C)) *sap130b ^pt35b/pt35b^* that introduced a premature stop codon in exon 6 of *sap130b* disrupting the N-terminus and eliminating the C-terminal region (**Figure 1C, Supplementary Tables S2, S3**). Using the *Tg(myl7:EGFP)* line, which labels the heart with green fluorescent protein, we found that 48% of the *MZsap130a;Tg(myl7:EGFP)* mutant embryos had the SV heart phenotype at 48hpf (**Figure 1F**). In contrast, only 17% of the *MZsap130b;Tg(myl7:EGFP)* mutant embryos had SVs by 48hpf (**Figure 1F**). We generated double mutants to further explore if *sap130a* and *sap130b* have any redundant functions (**Figure 2A, B**). We measured these adults from a double heterozygous in-cross and found that the *sap130a/b* double homozygous mutants are much smaller than their double heterozygous siblings (**Figure 2C**). *MZsap130a;sap130b^pt35b/+^* mutant in-crosses, revealed 39% of the embryos had SVs at 48hpf (**Figure 2D**). These observations suggest *sap130b, unlike sap130a,* is not required for zebrafish cardiogenesis.

**Figure 1:**
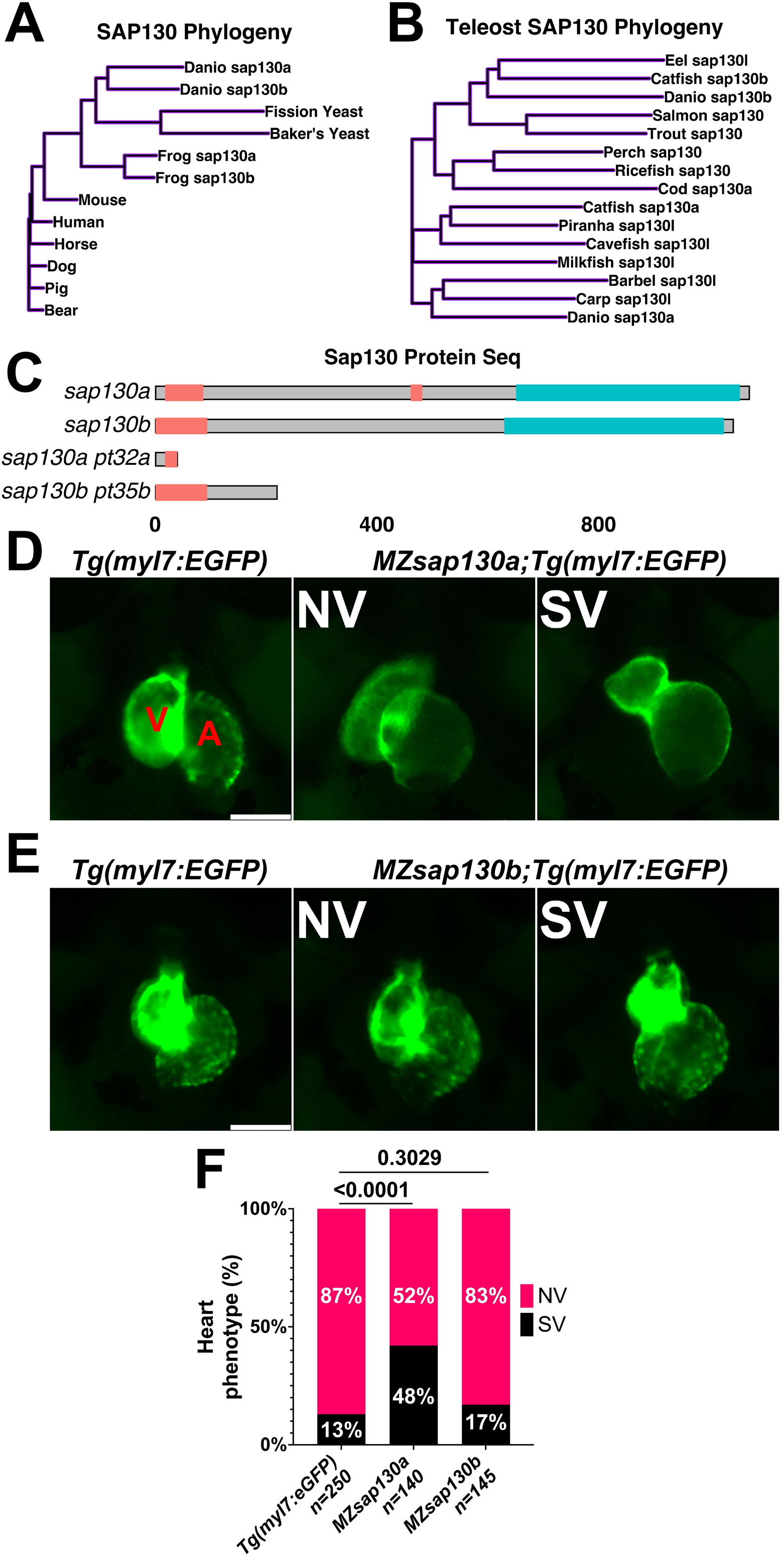
*sap130a* and *sap130b* have non-overlapping functions in the zebrafish heart. (**A, B**) A simple distance matrix phylogeny tree of Sap130a and Sap130b in broad or teleost specific contexts (**C**) Schematic of Sap130a and Sap130b protein sequences from the UniProt database highlighting the conserved regions and predicted mutant proteins. Unorganized sequence in pink, C-terminal conserved domain in blue, which contains the binding domain for SIN3A and HDAC1 (**D, E**) Representative images of *Tg(myl7:EGFP*), *MZsap130a;Tg(myl7:EGFP)* and *MZsap130b;Tg(myl7:EGFP)* mutant hearts at 48hpf (**F**) Graph of heart phenotype (%) proportions, pvals from fisher’s exact test. V and A are ventricle and atria, respectively. Scale bar: 100μm.

**Figure 2:**
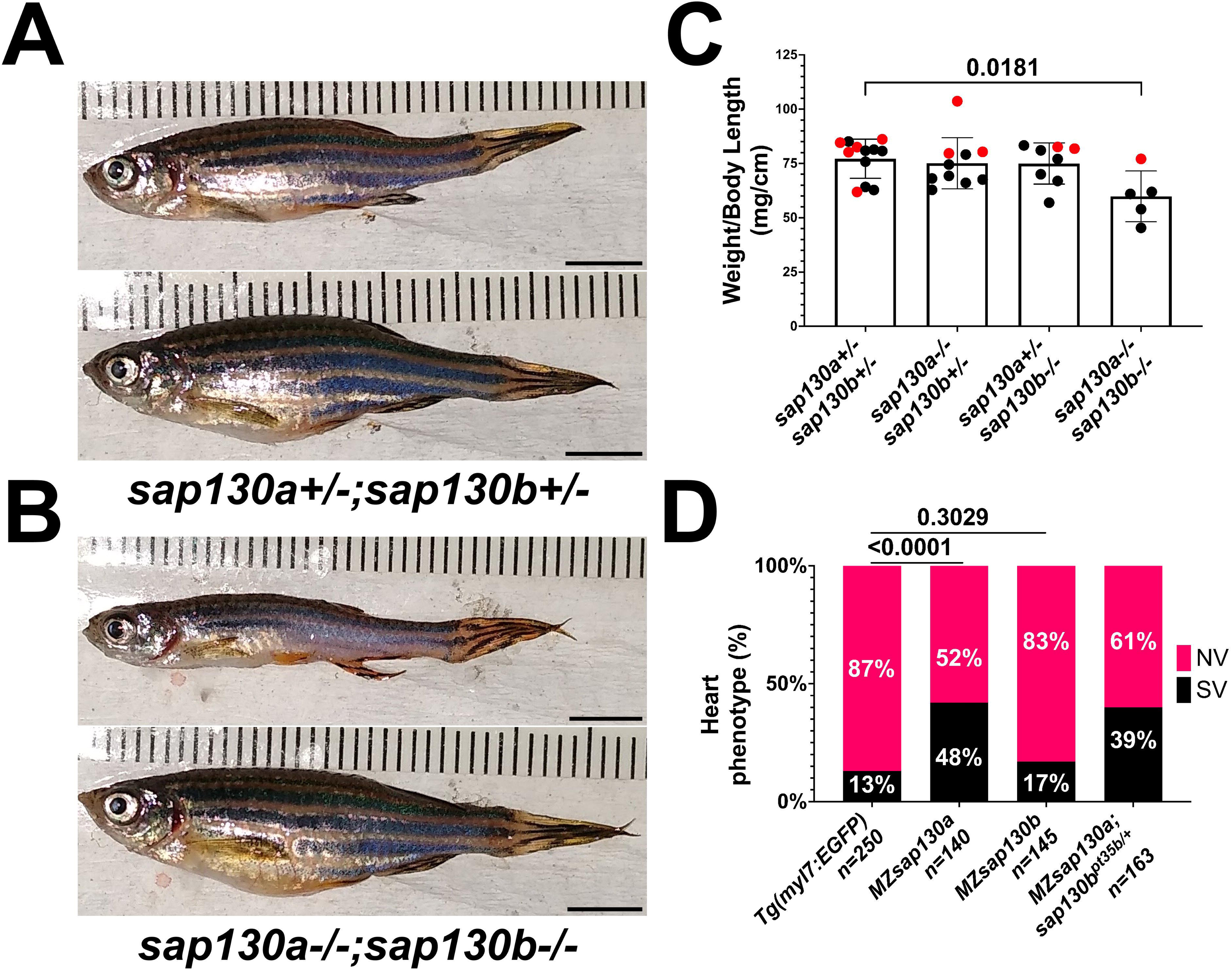
Double *sap130a; sap130b* mutants fail to thrive as adults. (**A**) *sap130a*;*sap130b* double heterozygous adults, male (top) and female (bottom). (**B**) *sap130a*;*sap130b* double homozygous adults, male (top) and female (bottom). (**C**) Graph quantifying weight to length ratio for adults from a *sap130a*;*sap130b* double heterozygous in-cross, pvals are for one-way ANOVA, error bars are standard error mean (SEM). Red points represent females and males in black. (**D**) Graph quantifying the heart phenotype at 48hpf proportions for *Tg(myl7:EGFP*), *MZsap130a;Tg(myl7:EGFP)*, *MZsap130b;Tg(myl7:EGFP*), and *MZsap130a;sap130b^pt35b/+^;Tg(myl7:EGFP)*, pvals are for fisher’s exact test. Scale bar: 5 mm

*sap130a* AUG start codon antisense-morpholino (MO) studies suggested the SVs arise from decreased ventricular CMs (7), but where or when CMs are lost was not explored. To determine if the SVs are due to decreased cardiac progenitors, we performed Whole Mount In Situ Hybridization (WISH) at 10 somite stage with *nkx2.5*, an early cardiac progenitor marker. We discovered no differences between *MZsap130a* and controls (**Figure 3A**). This suggests that the early cardiac progenitors were present in the *MZsap130a* embryos. To profile the chambers of the heart we performed WISH at 24hpf with myosin heavy chain 7 (*myh7*, ventricle) and myosin heavy chain 6 (*myh6*, atria). These probes did not reveal any difference between WT and mutant embryos, suggesting the First Heart Field is intact (**Figure 3B**). At 36hpf and 48hpf the atrial chamber showed no change, but the ventricle was smaller (**Figure 3C, Figure S2**). This phenotype was observed again when imaging the *MZsap130a;Tg(myl7:EGFP)* at 36hpf (**Figure 4A**). Many studies have detailed the second heart field accretion between 24 and 48hpf in zebrafish (38, 44, 50, 51). These SHF cells trail behind the heart tube and add to the ventricle continuously. There is speculation as to how many SHF cells are ventricular CMs, debated to contribute between 30-40% of the total ventricular CMs by 48hpf (49). The SV heart phenotype arising at 36hpf and the lack of changes seen in FHF markers suggest the SHF might be an influenced cell population where CMs are lost in *MZsap130a* mutants.

**Figure 3:**
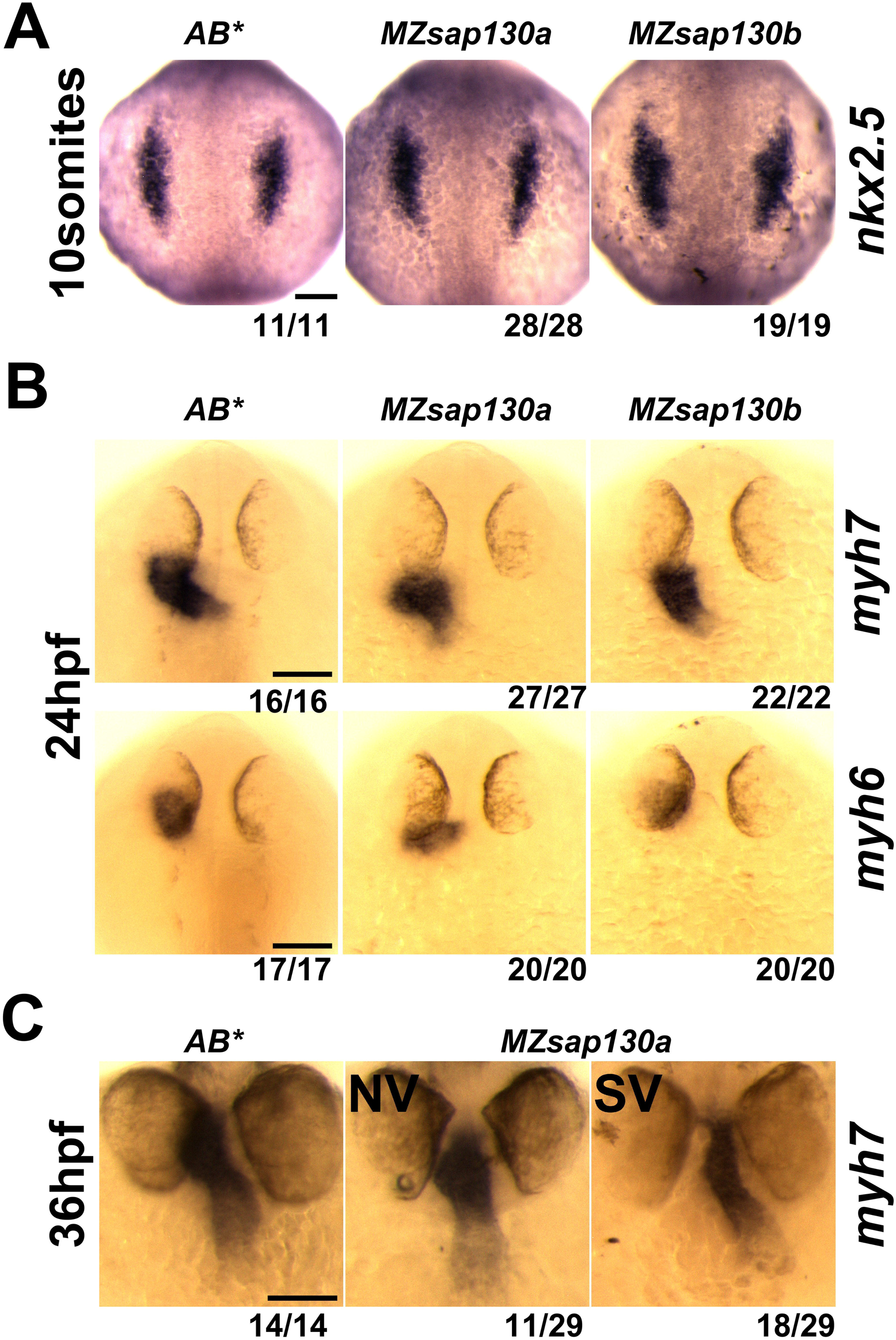
Cardiac gene expression in *MZsap130a* and *MZsap130b*. (**A**) WISH of *nkx2.5* at 10 somite stage for *AB\**, *MZsap130a* and *MZsap130b*. (**B**) WISH of *myh6* and *myh7* at 24hpf for *AB\**, *MZsap130a* and *MZsap130b*. (**C**) WISH of *myh7* at 36hpf in *AB\** and *MZsap130a*. Scale bar: 100μm

**Figure 4:**
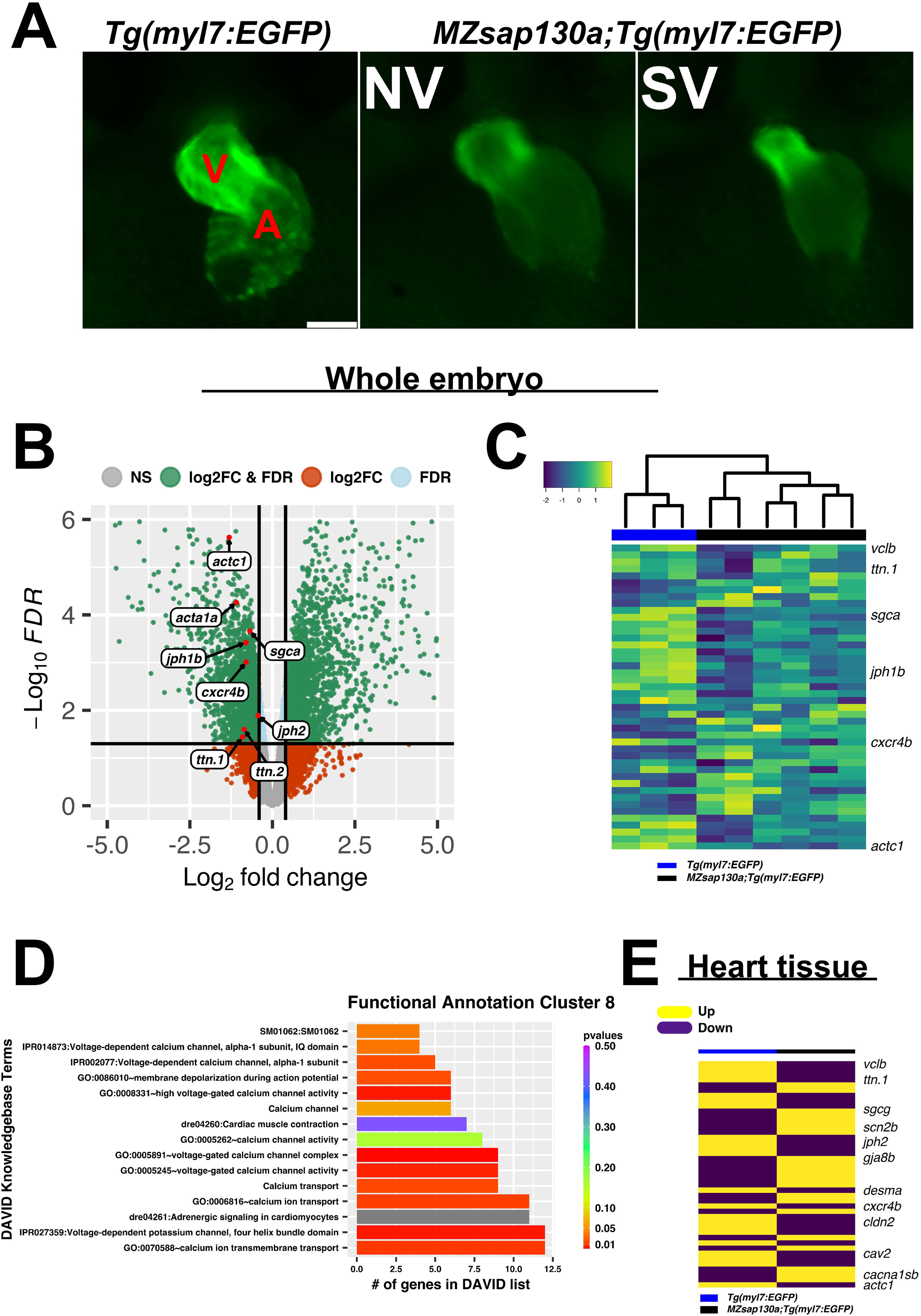
RNA-seq reveals cardiac sarcomere and calcium channels are altered in *MZsap130a* mutants. (**A**) Representative images of *Tg(myl7:EGFP*) and *MZsap130a;Tg(myl7:EGFP)* embryos collected for whole embryo RNAseq at 36hpf. (**B**) Volcano plot of 36hpf whole embryo RNA-seq data. (**C**) Heatmap of sarcomere and calcium channels genes (**Supplementary Table S6**) at 36hpf from whole embryos. (**D**) DAVID functional annotation cluster 8 from down-regulated genes in *MZsap130a*, showing DAVID calculated p-values. (**E**) Heatmap of 48hpf heart tissue RNAseq data for the same genes found in panel C. V and A are ventricle and atria, respectively. Scale bar: 100μm

### RNAseq reveals *sap130a* is involved in regulating cardiac sarcomere and calcium channel gene expression in zebrafish

Variants in genes encoding sarcomere proteins have been linked to CHDs (2, 77, 78, 79, 80, 81, 82, 83, 84). The Sin3A complex has been shown to regulate sarcomere specific genes like titins, troponins, and actins important for cardiac contraction (24). Since SAP130 has been shown to be part of the SIN3A complex, we reasoned that the phenotype may be caused by altered regulation of cardiac gene expression during development. A whole embryo RNA sequencing (RNAseq) experiment, separating the SV and “normal” (NV) siblings in the *MZsap130a* mutants, was performed at 36hpf. We first performed our analysis looking for differences in the wildtype, compared to NV and SV separately finding 2826 differentially expressed genes (DEGs) in common, with 812 unique DEGs for NV and 1979 for SV. Functional annotation of these gene groups revealed that NV and SV embryos are similar when compared to the wildtype transcriptome (**Supplementary Table S4**). Comparing the controls to all *MZsap130a* samples, we observed 5002 DEGs that included many cardiac specific transcripts. Among the DEGs we found sarcomere and calcium channel genes were dysregulated, suggesting CM function has changed in the *MZsap130a* embryos (**Figure 4B, C, Supplementary Tables S5, S6**). DAVID functional annotation of down regulated genes showed enrichment for cardiac contraction and adrenergic signaling in CMs, further suggesting a role for *sap130a* in CM function (**Figure 4D, Supplementary Table S5**). To confirm cardiac specific changes in these same transcripts, *MZsap130a* mutant hearts and controls were harvested at 48hpf and the transcriptome was profiled, showing similar cardiac gene expression changes (**Figure 4E, Supplementary Table S7**). We also profiled the *MZsap130b* transcriptome at 36hpf and found less gene expression changes (617 DEGs), without the same sarcomere changes seen in *MZsap130a* (**Figure S3A, B, C, D, Supplementary Table S8**). DAVID functional annotation of the 278 DEGs common between *MZsap130a* and *MZsap130b* mutants, belonged to heme binding and biosynthesis, oxygen binding, and iron binding KEGG pathways, suggesting involvement in hematopoiesis (**Supplementary Table S8**). The expression profile for these hematopoietic related genes was opposite in *MZsap130a* and *MZsap130b*, suggesting distinct functions during hematopoiesis (**Figure S3E, F, G**). These data suggest that *sap130a* and *sap130b* could be involved in hematopoiesis, consistent with *sin3aa*/*ab* gene knockdown studies showing strong hematopoietic defects (27).

Whole embryo and heart tissue *MZsap130a* RNA-seq data revealed sarcomere genes such as actins and myosins were dysregulated, indicating that sarcomere dysfunction could be part for the *MZsap130a* mutant phenotype. These data also showed down regulation of CM cell communication genes such as *cxcr4b* and *gja3*, also resulting in small ventricle phenotypes when mutated in zebrafish (85, 86, 87, 88, 89). Changes were also found in calcium channel (*cacna1sb*, *cacng7a*, *cacnb1*, *cacna1bb*) and sodium channel (*scn4aa*/*ab*, *scn2b*) genes, known to be important to CM functional activity (90, 91, 92). Furthermore, transcriptome analysis revealed that *MZsap130a* mutants showed dysregulation of a wide range of genes critical for cardiac maturation and function. These include genes associated with fatty acid metabolism (*ppt2*), glycogen metabolism (*ugp2a*, *phka2*), and mitochondria (*slc25a44a*, *slc25a42*, *mtrf1*, *mrpl58*) found down regulated in *MZsap130a* mutants in whole embryos at 36hpf and specifically in the heart at 48hpf (**Figure S4C, F**). Collectively, *MZsap130a* mutants show changes in sarcomere, cell communication and metabolism associated genes, all integral parts of CM maturation.

### Sap130a regulates cardiac function

Global loss of *sap130a* showed downregulation of cardiac and skeletal sarcomere genes such as *actc1*, *ttn.1*, and *ttn.2* (**Figure 4B-E**). This suggested that cardiac function could be diminished in *MZsap130a* mutants. The DAVID functional annotation tool revealed enrichment for Cardiac Contraction genes that were decreased in the *MZsap130a* mutant embryos (**Figure 4D**). To determine ventricle chamber function in mutants, confocal light sheet microscopy was used to record live cardiac contractions at 48hpf. These recordings provided us with multiple frames of diastole and systole for chamber volume estimation (**Figure 5A-C, and Supplementary Movie S1, S2, S3, S4, S5**). Volume estimations were used to calculate the cardiac parameters Total Stroke Volume (TSV), and Cardiac Output (CO) (75). The light sheet data revealed that all *MZsap130a* mutants had deficits in CO, TSV, fractional shortening, and ejection fraction. In comparison, the *MZsap130b* mutant heart revealed no significant difference from WT hearts, both in TSV and CO (**Figure 5**). Our heart tissue RNA-seq found cardiac contraction genes *myh7*, *actc1*, *ttn.1*, *ttn.2*, *scn4ab*, and *cacna1sb* were dysregulated in *MZsap130a* mutants, supporting the contraction deficits at 48hpf (**Figure S4D**). These data show that *sap130a* has a role in zebrafish cardiac sarcomere regulation.

**Figure 5:**
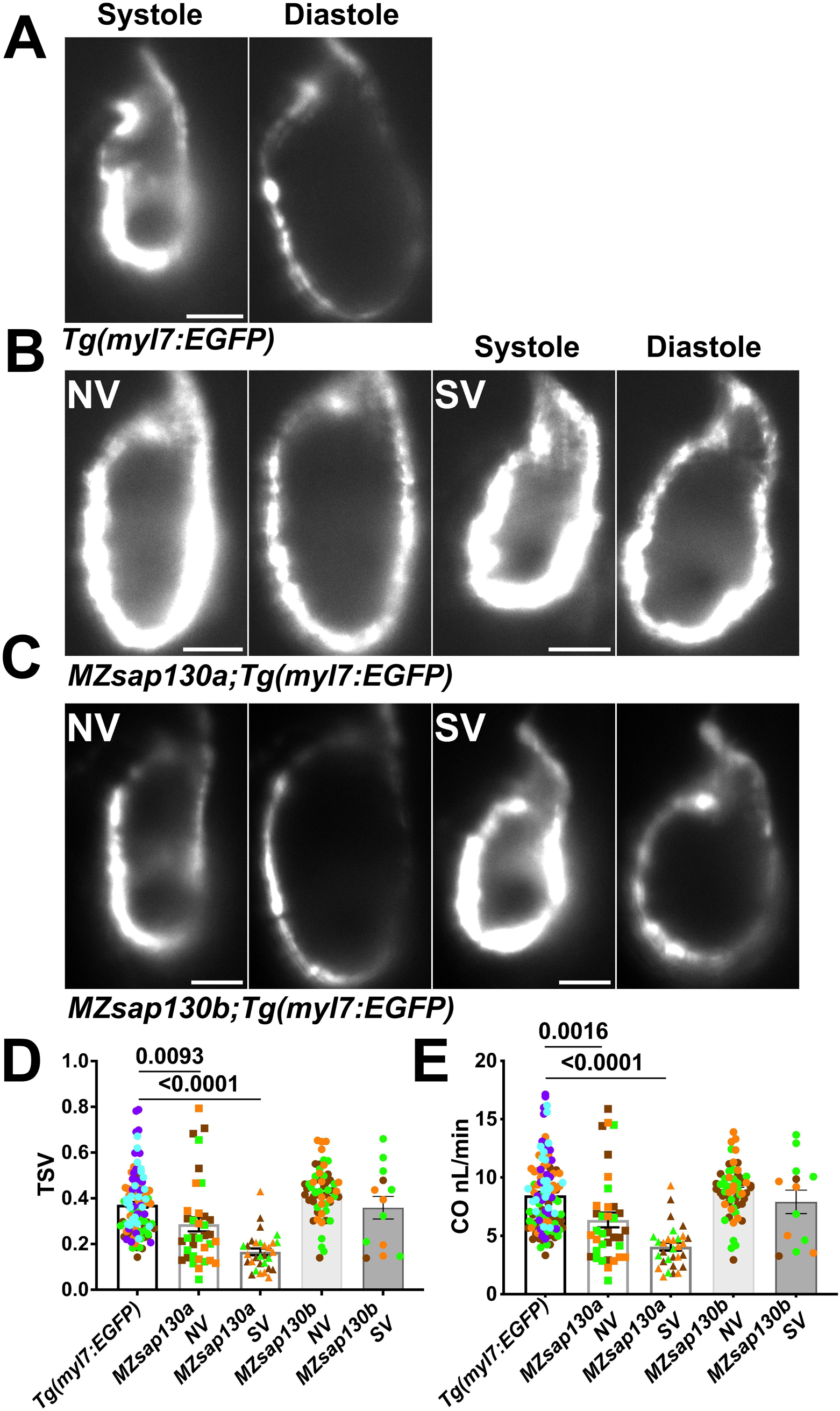
*MZsap130a* embryos show cardiac functional deficits. (**A, B, C**) Shows systole and diastole frames from recordings of live ventricular contractions in *Tg(myl7:EGFP*), *MZsap130a;Tg(myl7:EGFP)* and *MZsap130b;Tg(myl7:EGFP*) at 48hpf. (**D, E**) Quantified cardiac parameters total stroke volume (TSV) and cardiac output (CO), pvals from one-way ANOVA, error bars are SEM. Each point represents individual ventricle and color coded for 3+ experiments. For *Tg(myl7:EGFP)*, n=115; *MZsap130a* NV, n=36; *MZsap130a* SV, n=30; *MZsap130b* NV, n=57; *MZsap130b* SV, n=13. Scale bar: 20μm

### *MZsap130a* mutants have longer outflow tracts

The earliest observation of smaller ventricles in *MZsap130a* mutants was at 36hpf, a stage when SHF cells are migrating into the ventricle. Extensive studies have reported the contribution of SHF cells to the ventricle during this time (19, 44, 46, 49, 50, 51). Our RNA-seq data showed that SHF progenitor markers *ltbp3*, *mef2cb* and *isl1, isl2a/b* were decreased (**Figure S4B, E**). These genes are known to label SHF progenitors at the arterial and venous poles. This suggested that the SHF in the *MZsap130a* mutants was affected such that insufficient CMs contribute to the ventricle by 48hpf. To determine if this occurs, we performed lineage tracing experiments using *Tg(nkx2.5:kaede)* embryos (45). In this transgenic line, the FHF cells can be permanently labeled at 24hpf, photo-converting only the heart tube. Next, we image at 48hpf to determine the addition of green cells to the ventricle (**Figure 6A, Figure S6**). We lineage traced the SHF with *MZsap130a;Tg(nkx2.5:kaede)* embryos and revealed that the SVs acquire less SHF (green area) compared to the wildtype and *MZsap130a* mutant siblings that develop normal ventricles (**Figure 6B, C, D**). Moreover, the OFTs in the *MZsap130a* mutants were longer at 48hpf in some embryos with SVs (**Figure S7A, B**). The longer OFTs were much more pronounced at 72hpf, and every SV heart had a longer OFT (**Figure 6E, F, Figure S7C, D**). This suggested that the lost ventricular CMs contributed to OFT cells instead and was further evidenced at adult stages. *MZsap130a;Tg(myl7:EGFP)* embryos were scored at 48hpf for ventricle size and reared separately into adulthood. Brightfield images of heart extractions revealed a larger bulbus arteriosus (BA) area, the adult structure derived from the OFT, and less ventricular area (**Figure 7**). This suggests *sap130a* is involved in SHF cell fate decisions between ventricular CMs and OFT cells.

**Figure 6:**
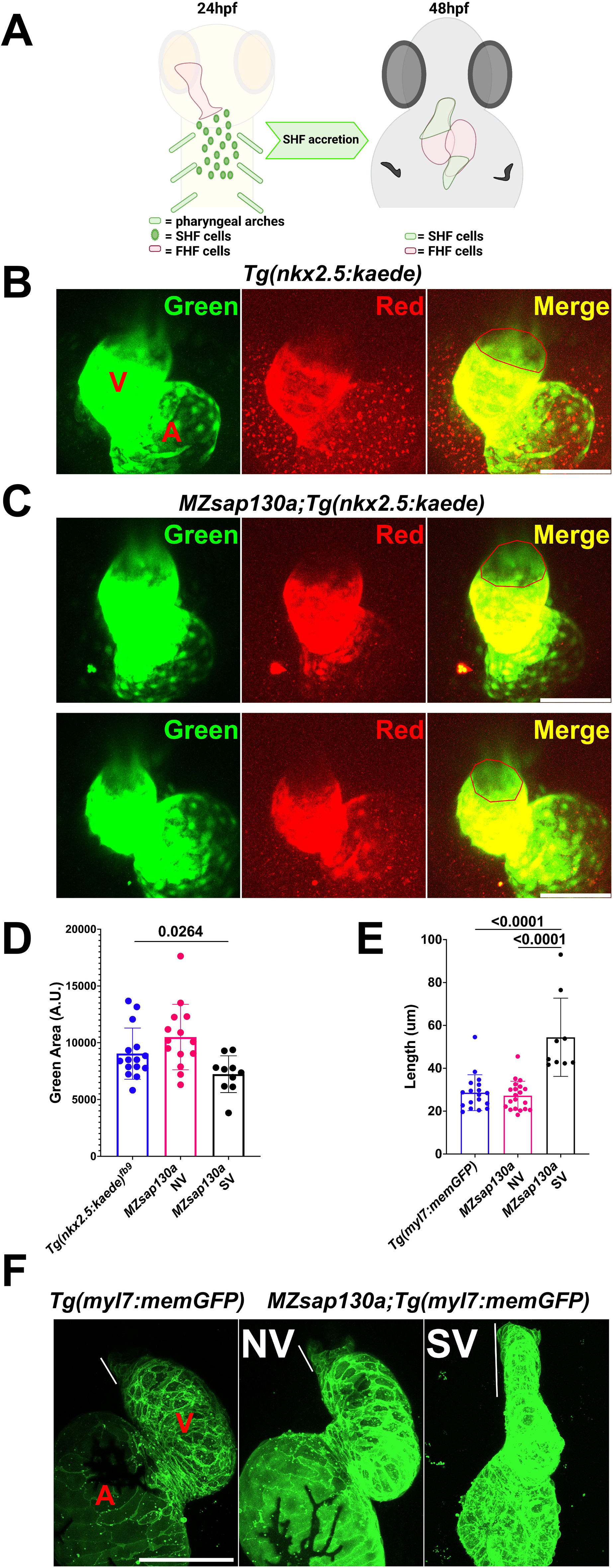
Lineage tracing reveal changes to SHF and OFT in *MZsap130a*. (**A**) Diagram showing how the FHF heart tube at 24hpf was photoconverted to red, leaving SHF progenitors unlabeled in green and imaging at 48hpf. (**B, C**) Confocal imagines of *Tg(nkx2.5:kaede)* and *MZsap130a;Tg(nkx2.5:kaede)* at 48hpf with the heart tube being photoconverted at 24hpf, the red outlined region represents area measurements collected. (**D**) Quantified SHF (green area) accreted by 48hpf, pval is from a one ANOVA, error bars are SEM. Each point represents a single embryo, *Tg(nkx2.5:kaede)*, n=15; *MZsap130a;Tg(nkx2.5:kaede)* NV, n=14; *MZsap130a;Tg(nkx2.5:kaede)* SV, n=10. (**E**) Quantified OFT lengths at 72hpf, pval is from a one ANOVA, error bars are SEM. Each point represents a single embryo, *Tg(myl7:memGFP)*, n=18; *MZsap130a;Tg(myl7:memGFP)* NV, n=20; *MZsap130a;Tg(myl7:memGFP)* SV, n=9 (**F**) Representative images of *Tg(myl7:memGFP)* and *MZsap130a;Tg(myl7:memGFP)*, white lines demarcate OFT length, pvals from one-way ANOVA, error bars are SEM. V and A are ventricle and atria, respectively. Scale bar: 100μm

**Figure 7:**
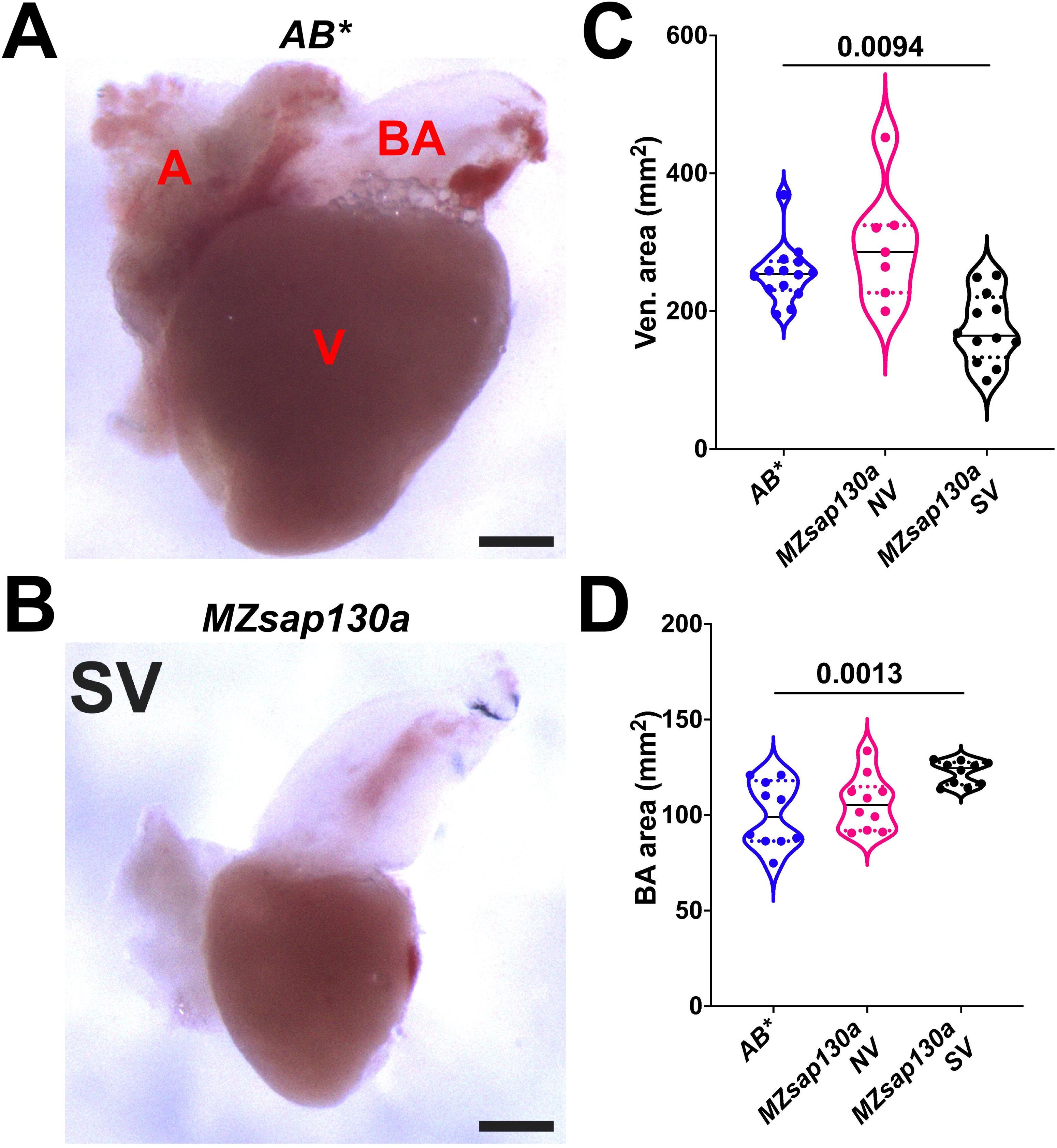
Adult *MZsap130a* hearts have large bulbus arteriosus. (**A, B**) *AB\** and *MZsap130a* adult SV hearts extracted at approximately 4-6 months post fertilization. (**C**) Quantification of ventricle area of unfixed hearts, pval is from a one ANOVA solid black bars are mean, dotted lines represent up and lower 25^th^ percentiles. Each point represents a single heart, AB*, n =14; *MZsap130a* NV, n=7; *MZsap130a* SV, n=12. (D) Quantification of BA area. of unfixed hearts, pval is from a one-way ANOVA solid black bars are mean, dotted lines represent up and lower 25^th^ percentiles. Each point represents a single heart, n=10 for all groups. V, A, and BA are ventricle, atria, and bulbus arteriosus respectively, Scale bar: 200μm

### *sap130a* genetically interacts with *hdac1* during SHF accretion

Zebrafish *hdac1* is required for ventricle formation (18, 19). We explored the potential interaction of Sap130a and Hdac1 by analyzing heart development in *MZsap130a;hdac1^+/b382^* embryos. While *hdac1* homozygous mutants develop cardiac defects, heterozygous mutants are viable and generally, ventricle size appear normal. In contrast, an increase in SV phenotype was noted in *MZsap130a;hdac1^+/b382^* suggesting *MZsap130a* mutants are sensitized to *hdac1* gene dosage (**Figure 8A, C**). These data revealed an association between *hdac1* heterozygous status and ventricle size only when in a *MZsap130a* background (**Figure 8C**). This suggests that *sap130a* and *hdac1* genetically interact in zebrafish and that these proteins function in the same complex.

**Figure 8:**
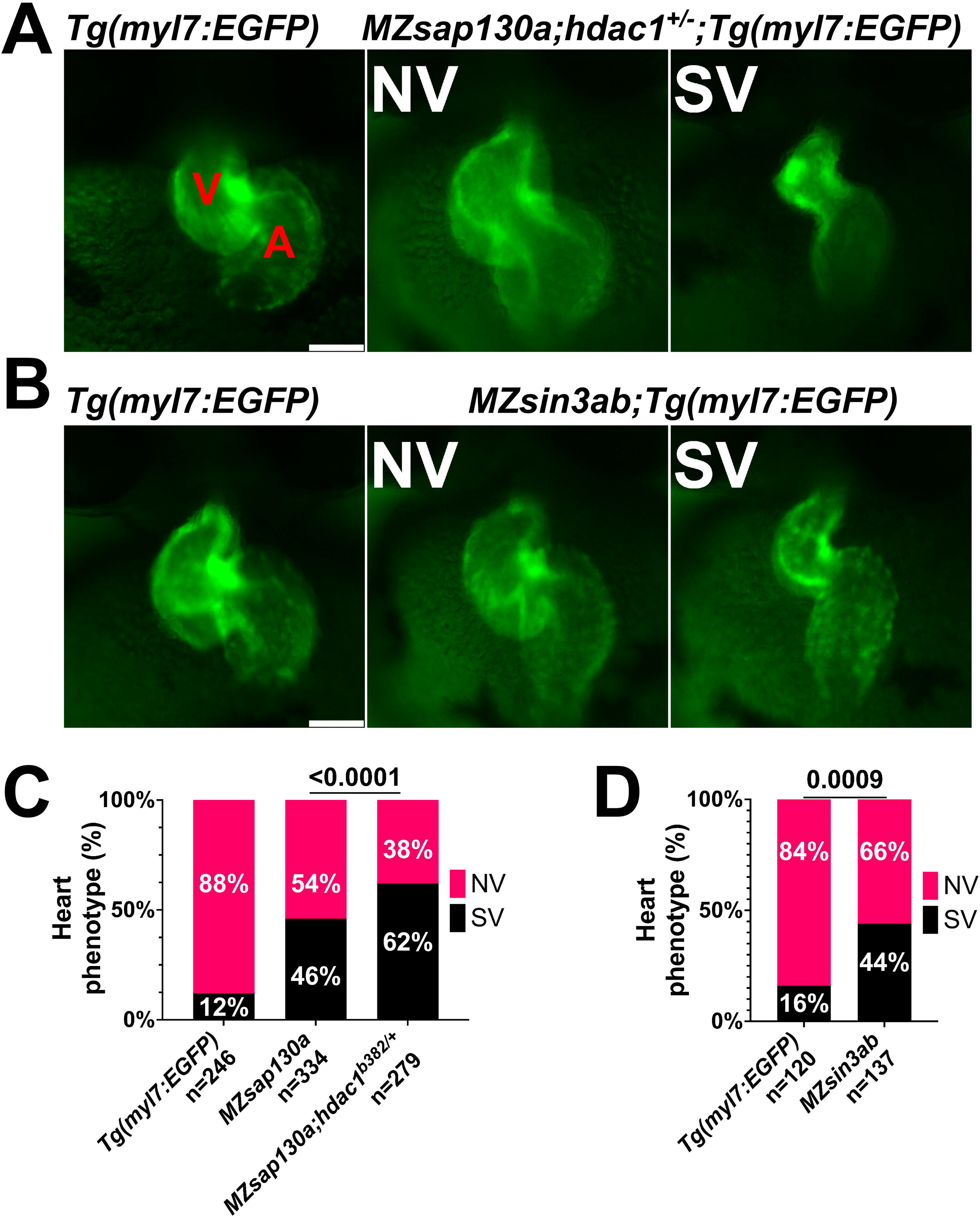
Genetic studies reveal the importance of the *sin3ab*, *sap130a* and *hdac1* in heart development. (**A**) Representative image of *Tg(myl7:EGFP)* and *MZsap130a;hdac1^-/+^;Tg(myl7:EGFP)* NV and SVs at 48hpf. (**B**) Representative image of *Tg(myl7:EGFP)* and *MZsin3ab*;*Tg(myl7:EGFP)* NV and SVs at 48hpf. (**C**) Quantification of heart phenotype proportions in *Tg(myl7:EGFP)*, *MZsap130a;Tg(myl7:EGFP)*, and *MZsap130a;hdac1^-/+^;Tg(myl7:EGFP)*. The p-values are fisher’s exact test. (**D**) Quantification of heart phenotype proportions in *Tg(myl7:EGFP)* and *MZsin3ab;Tg(myl7:EGFP).* The p-values are fisher’s exact test. V and A are ventricle and atria, respectively. Scale bar: 100μm

Both *MZsap130a* whole embryo and heart specific RNA-seq datasets revealed sarcomere genes to be down and cell cycle genes to be up regulated, similar to SIN3A knock-out and knock-down studies (**Figure S4A**) (23, 24, 25, 93). For example, the cell cycle genes *vrk1*, *rb1*, *e2f7*, and *cdkn1a* are found upregulated in *MZsap130a* mutants, while we do not find evidence of expanded progenitor populations. These similarities in up and down DEGs point to the possibility that *sap130a* associates with *sin3aa* or *sin3ab* in zebrafish, similar to mammals. To further explore the importance of SIN3A in heart development, we generated *MZsin3ab* mutants using CRISPR/Cas9. We generated a *sin3ab pt36a* allele (5bp del, 2bp insertion, 1bp substitution (T>C)) that disrupted amino acids 862-867 (**Figure S8A**). Only maternal and zygotic loss of *sin3ab* was sufficient to evoke the SV phenotype in 44% at 48hpf (**Figure 8B, D**). It is not surprising that the penetrance of the phenotype in *MZsin3ab* was also incomplete since *sin3aa* and *sin3b* could compensate for the loss of *sin3ab*. These data suggest that *sin3ab* is involved in ventricular development in zebrafish, a phenotype that is similar to *sap130a* and *hdac1* mutants.

## Discussion

In this study, we have revealed a role for *sap130a* in zebrafish cardiogenesis. We describe a null allele of *sap130a*, resulting in small ventricles through the delay and failure of SHF cells to migrate into the ventricle. Without *sap130a*, some of the SHF progenitors permanently become OFT cells. Transcriptome profiling of the *MZsap130a* embryos at 36hpf and hearts at 48hpf revealed that expression of sarcomere, cell communication, and metabolism genes were dysregulated. This suggest that the CMs fail to terminally differentiate and properly function.

Our study reveals the consequence of disrupting members of the SIN3A complex, resulting in improper heart development. In the *MZsap130a* mutants, the main phenotype is a small ventricle leading to larger OFT and bulbus arteriosus in adults. Developmentally this arises from the failure of SHF progenitors to migrate into the growing ventricle. We come to this conclusion because the WISH data for *nkx2.5 myh7* showed no changes prior to the 24hpf, indicating the FHF is intact. The phenotype arising at 36hpf is in line with observations showing the addition of SHF cells between 24-48hpf and with our lineage tracing experiments (38, 44, 49, 50, 51). In the *Ohia* mouse mutant, the combination of *PCDHA9* and a *SAP130* mutations caused an HLHS etiology influencing the FHF structures. The prominent phenotype included a hypoplastic left ventricle and valve abnormalities in 11% of mouse embryos. In the zebrafish, the *sap130a* mutation is predicted to be a null mutant producing a hypoplastic ventricle in 48% of embryos. The difference seen between the mouse and zebrafish can be explained by the difference in the number of ventricle chambers, the specialized development of the mammalian OFT, and the changes seen during the evolution of this specialized pump (32, 33, 35, 36). A recent study of *SAP130* pig CRISPR mutants show tricuspid dysplasia and atresia, highlighting the complex role of Sap130 in heart development across different species (8).

Unlike mammals, where SAP130 is critical for survival, loss of *sap130a* produced embryos that had an incomplete penetrance of cardiac defects, while *sap130b* were normal. The changes noted in sarcomere transcripts in the *MZsap130a* mutants resembles genes that SIN3A has been shown to target in skeletal muscle (24, 25, 94). In this study, cell cycle genes were upregulated upon knock-down of SIN3A in C2C12 myotubes, and interestingly we see similar upregulation in our *MZsap130a* mutants (24). One conclusion supports the notion that SIN3A activity is impaired in the absence of SAP130. This supports the idea that *sap130a* and *sin3aa*/*ab* in zebrafish could work in complex, as they do in mouse and human cell lines. This implicates *sap130a* as playing a critical role in regulating muscle cell differentiation within the SIN3A complex. In zebrafish, genetic interaction studies could be done for *sap130a* and *sin3aa*/*ab* as well as injection of antisense morpholinos targeting that same combination of genes. Although we showed *MZsin3ab* mutants have SVs, suggesting that the *sin3a* genes are important for heart development, we have not yet tested *sin3ab* and *sap130a* genetic interaction. Further studies are required to determine if *sap130a* associates with *sin3aa*/*ab*.

The DAVID functional annotation tool revealed many changes between the wildtype and *MZsap130a* mutants, including those in adrenergic signaling in CMs. Adrenergic signaling has been shown to be involved in the metabolic switch from glycolysis to the Krebs cycle, a process that is required for CM maturation. Adrenergic deficient mice showed poor CM mitochondrial function and physiology (95, 96). Interestingly, *MZsap130a* mutants show a decrease in adrenergic signaling genes *bcl2b*, *camk2d2*, *adcy3a*, *actc2*, and *gna15.1*, implicating SAP130 is involved in mitochondrial biology (97). Evidence for adrenergic signaling being epigenetically regulated and interacting with troponin T of the sarcomere could further suggest *sap130a*’s involvement in this metabolic process required for CM maturation (98, 99, 100). Further annotation of *MZsap130a* mutant hearts identified the same genes involved in metabolism and mitochondria to be dysregulated in the whole embryo. This suggests that the maturation of the CM mitochondria has not recovered by 48hpf. A defect in mitochondrial metabolism has been observed in the heart tissue and cardiomyocytes derived from the *Ohia* mutant mice, and in HLHS patient heart tissue and iPSC derived cardiomyocytes (97). In both the HLHS mouse and human cardiomyocytes, defects are also noted in cardiomyocyte differentiation and maturation, and this is accompanied by poor contractile function. Our findings in this study agree with human and mouse data (7, 97). Of further note, mitochondrial respiration and sarcomere assembly deficits were also observed in zebrafish with mutations in *rbfox1l* and *rbfox2*, a gene family that have been associated with HLHS (101). Double *rbfox* mutants show impaired myocardial contractility and malformed mitochondria. Thus, the phenotypes observed in the *rbfox* double mutants and *MZsap130a* are the result of changes in genes that are important for the maturation of CMs and their function. Understanding the connections between the metabolism and CM maturation will allow for more detailed organoid model systems, as these studies usually lack terminally differentiated CMs (102).

The catalytic unit of the SIN3A complex is comprised of class I HDACs, which deacetylate lysine residues to alter gene expression or protein function. The *hdac1 b382* allele, used for the genetic interaction studies, is not the only *hdac1* zebrafish mutant shown to have cardiac defects (22). Another *hdac1* allele nl18, where a single nucleotide polymorphism disrupts the protein from exon7, has shown less ventricular CMs by 36hpf. This is also the earliest timepoint we report a difference in *MZsap130a* mutants. In the *hdac1/cardiac really gone* mutant, SHF cells do not proliferate, and transplantation studies revealed cell autonomous and nonautonomous functions in ventricular CMs (19). We did not observe a loss of CMs in the SHF, but a change from ventricular CMs to OFT cells in *MZsap130a* mutants. This suggests *sap130a* also has non-overlapping functions with hdac1 during cardiogenesis as has been suggested from in vitro studies (9).

The SIN3A/HDAC complex is known for histone deacetylation and has been shown to both inhibit and activate transcription events (10, 12, 103). These deacetylation events are sometimes paired with methylation events to regulate transcription (104, 105, 106, 107). One example of this is the SIN3A/HDAC complex and SMYD2 methylation of histone H3K36, which together repress cell proliferation in mouse fibroblasts (104). Another function of this SMYD2 protein is to methylate the cytoplasmic protein HSP90aa at K616 to form a complex to stabilize titin in the I-band of the sarcomere. This *smyd2*/*hsp90aa*/*ttn* support structure is needed when titin is put under mechanical stress and without it the sarcomeres are disorganized, particularly in the Z-disk and I-band structures (108, 109, 110). This structural support is a component of maturing sarcomeres, shown to be present in the skeletal and cardiac muscle tissues of mice, rat, chick, and zebrafish (108, 109, 110). Interestingly, *MZsap130a* RNA-seq data shows down regulation of *smyd2*/*4*, *hsp90aa*, and *ttn.1* and *ttn.2*. In the case of *MZsap130a* mutants, CM maturation is disrupted due to lack of proper sarcomere gene expression. The SIN3A/HDAC complex has been shown to be important in cell fate decisions and to be involved in myocyte maturation and sarcomere biology (13, 23, 24, 111). This work demonstrates that the Sap130/SIN3A/HDAC complex is involved in zebrafish cardiogenesis and allows us to study mutations in this complex usually lethal in mammals. These data build upon previous studies in zebrafish *hdac1* and reiterates the importance of context specific components for the SIN3A complex during cardiogenesis.

## Acknowledgements

RNA-seq data were aligned with CLC genomics Workbench Version 20 (QIAGEN), licensed through the Molecular Biology Information Service of the Health Sciences Library System, University of Pittsburgh. We are grateful to members of the Daniel Zuppo, Donghun Shin, Elizabeth Rochon, and Neil Hukriede for experimental suggestions. Tim Feinstein, who built the light sheet scope and mounting system. Manush Saydmohammed and Shoulin Li for initial work on *MZsap130a* mutant.

## Conflict of interest statement

All authors report that they have no conflict of interest with regards to this research.

## Author contributions

R.A.D, C.L. and M.T. designed research; R.A.D, R.F.R, D.F.J, and JS performed research; R.A.D and M.T. analyzed data. C.L., S.C.W. and M.T. responsible for funding. R.A.D wrote the manuscript. All authors edited the manuscript.

## Funding

This research was supported by funding from the National Institutes of Health R01HL142788 to M.T. and C.L., and R01HL16398 to M.T. In addition, support from the University of Pittsburgh Center for Research Computing, RRID:SCR_022735, and for the HTC cluster, which is supported by NIH award number S10OD028483. Genomics resources was provided through the University of Pittsburgh HSCRF Genomics Research Core, RRID: SCR_018301, for the RNA-seq experiments.

## Data Resource

The RNA-seq data files are available under the accession number: GSE228451

**Supplemental Information** Available as additional pdf document

---

**Supplementary Movie S1** show *Tg(myl7:EGFP)* ventricle at 48hpf

**Supplementary Movie S2** show *MZsap130a;Tg(myl7:EGFP)* with normal ventricle at 48hpf

**Supplementary Movie S3** show *MZsap130a;Tg(myl7:EGFP)* with small ventricle at 48hpf

**Supplementary Movie S4** show *MZsap130b;Tg(myl7:EGFP)* with normal ventricle at 48hpf

**Supplementary Movie S5** show *MZsap130b:Tg(myl7:EGFP)* with small ventricle at 48hpf

**Supplementary Table S1** Genotyping primers

**Supplementary Table S2** gRNA sequences

**Supplementary Table S3** Predicted amino acid sequence of sap130b and sin3ab mutation

**Supplementary Table S4** edgeR and DAVID results of WT vs *MZsap130a*NV and *MZsap130a*SV embryos at 36hpf

**Supplementary Table S5** edgeR and DAVID results of WT vs all *MZsap130a* embryos at 36hpf

**Supplementary Table S6** Log CMP of normalized counts for Figure 4C heatmap

**Supplementary Table S7** edgeR Likely Ratio Test of WT vs *MZsap130a* hearts at 48hpf

**Supplementary Table S8** edgeR and DAVID results of WT vs all *MZsap130b* embryos at 36hpf

## Notes

### Competing Interest Statement

The authors have declared no competing interest.

