## Supplemental Figures for "Sin3a Associated Protein 130kDa, sap130, plays an evolutionary conserved role in zebrafish heart development"

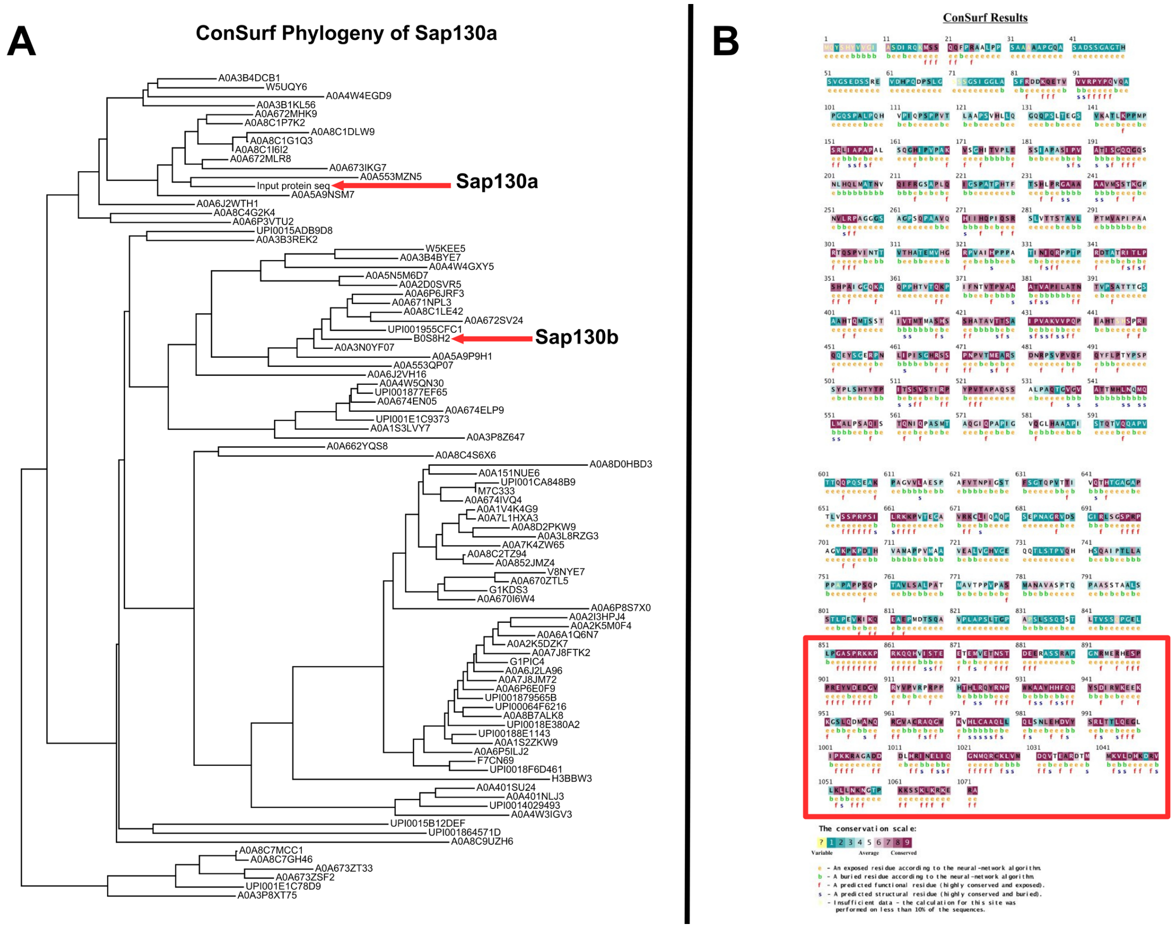


**FigS1: ConSurf output from sap130a protein sequence**

(**A**) A phylogenetic tree with UniProt IDs made using ConSurf showing 145 unique sequences for zebrafish sap130a. (**B**) The ConSurf output for protein residue classification, Darker red meaning more likely conserved. Red Box outlines the conserved C-terminal domain with many dark red amino acids.

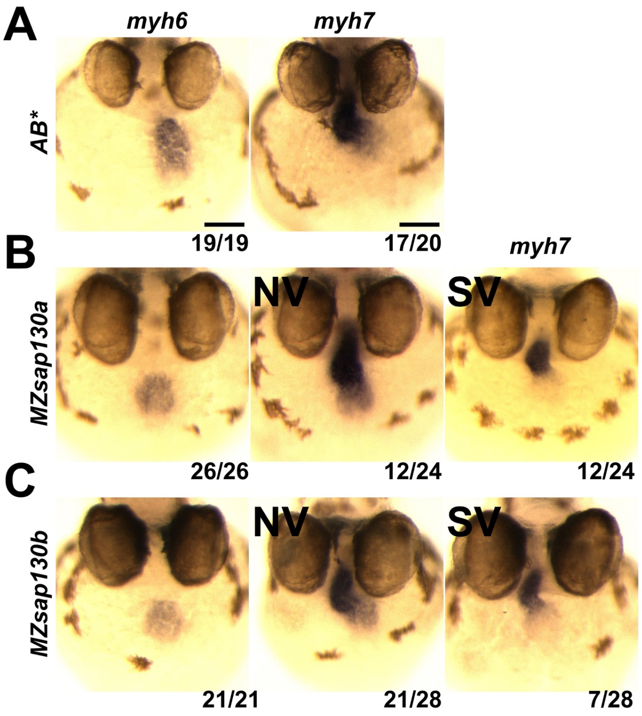


**FigS2: WISH at 48hpf of *MZsap130a* mutants have SVs**
(**A, B, C**) WISH of 48hpf embryos for *myh6* (atria) and *myh7* (ventricle) of AB* and *MZsap130a* and *MZsap130b* mutants. Scale bar 100µm

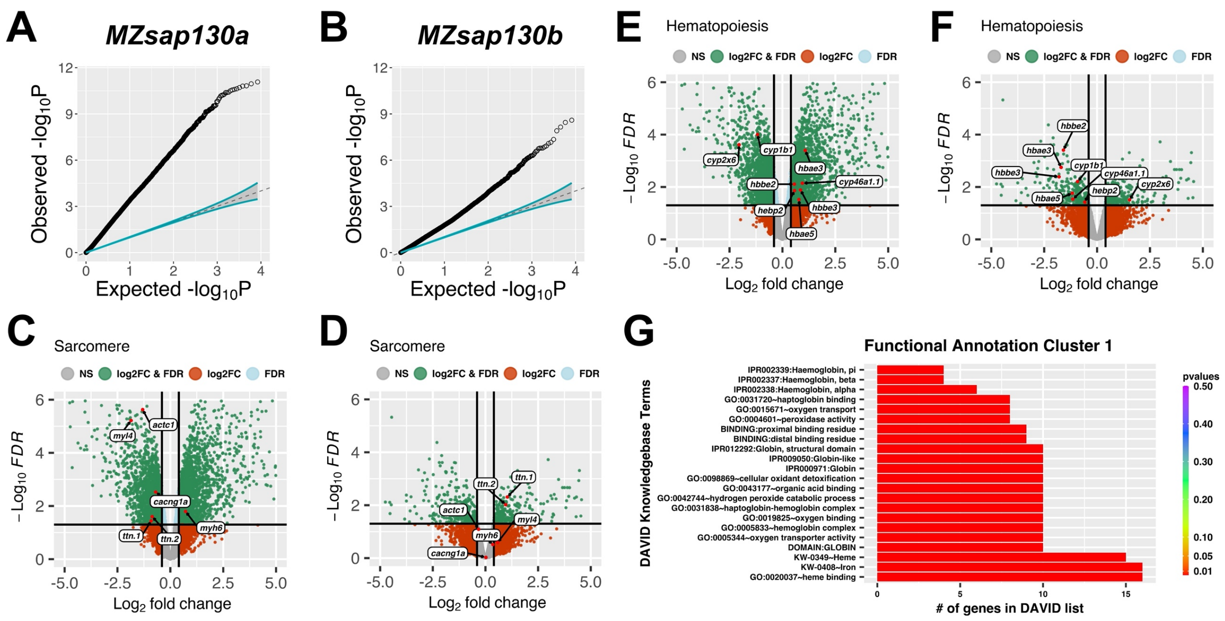


**FigS3: *MZsap130a* and *MZsap130b* show different gene expression profiles in zebrafish**
(**A, B**) Show the expected and observed –log10(pvals) for *MZsap130a* and *MZsap130b* mutants 36hpf whole embryo RNAseq. (**C, D**) Volcano plots showing changes in sarcomere genes in *MZsap130a* and *MZsap130b*. (**E ,F**) Volcano plots showing DEGs that overlap between *MZsap130a* and *MZsap130b*, involved in hematopoiesis. (**G**) Shows the DAVID annotation cluster 1 of the overlap genes revealing hematopoietic gene groups.

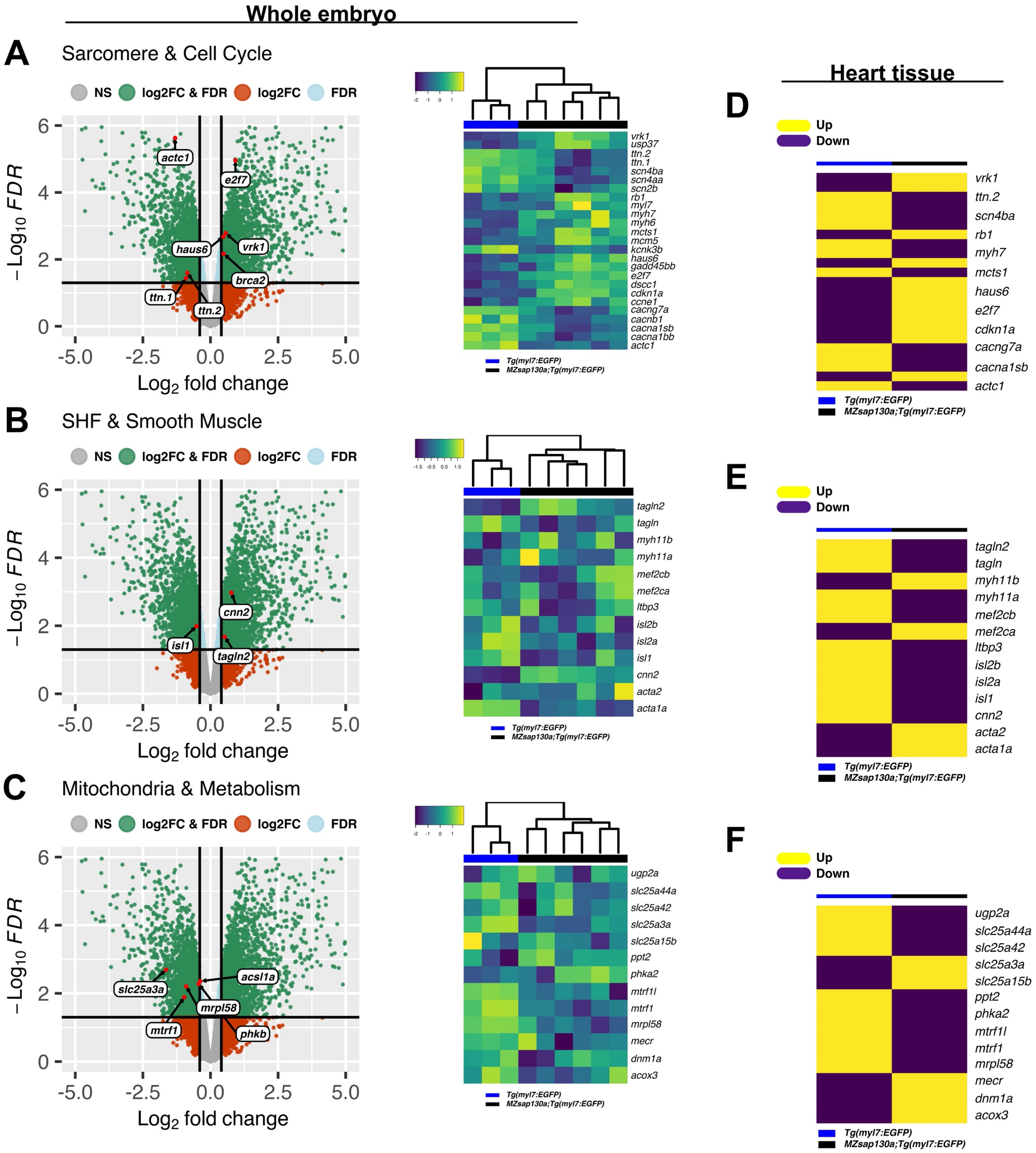


**FigS4: RNA-seq reveals DEGs and pathways in *MZsap130a* mutants**

(**A, B, C**) Volcano plots and heatmaps of genes associated with sarcomere and cell cycle, the SHF and smooth muscle, and metabolism genes from whole embryo RNAseq at 36hpf. (**D, E, F**) Heatmaps of the same genes in panels A, B and C, but for 48hpf heart tissue RNAseq. For 36hpf data an ANOVA-like analysis was used and for 48hpf heart tissue data we used a likelihood-ration test all in edgeR.

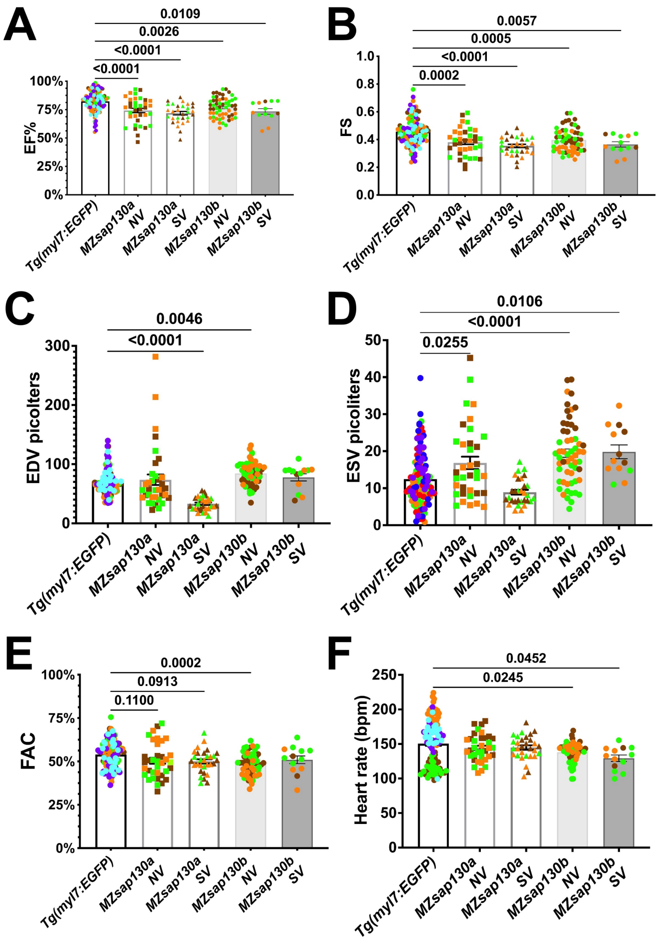


**FigS5: Cardiac parameters measured in *MZsap130* mutants**

(**A, B, C, D, E, F**) Shows the cardiac parameters measured ejection fraction (EF), fractional shortening (FS), end-diastolic volume and end-systolic volumes (EDV and ESV), fractional area change (FAC) and heart rate. Each point represents individual ventricle and color coded for 3+ experiments. For *Tg(myl7:EGFP)*, n=115; *MZsap130a* NV, n=36; *MZsap130a* SV, n=30; *MZsap130b* NV, n=57; *MZsap130b* SV, n=13.

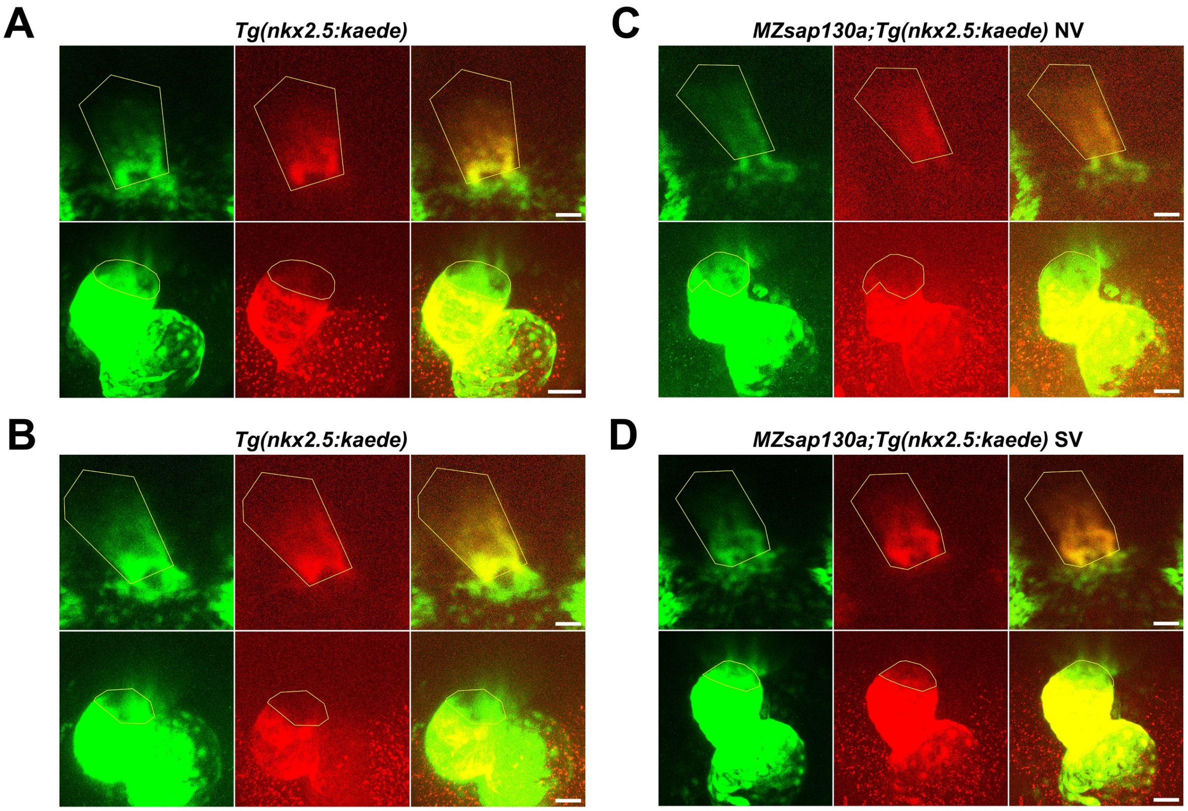


**FigS6: 24hpf heart tube photoconversion and 48hpf imaging and measuring examples**
(**A, B**) Shows examples of *Tg(nkx2.5:kaede)* being photoconverted at 24hpf and imaged at 48hpf. (**C, D**) Reveals another example of NV and SV *MZsap130a;Tg(nkx2.5:kaede)* mutants photoconverted at 24hpf and imaged at 48hpf. Yellow lines outline photoconverted region and the green area measured as SHF. *Tg(nkx2.5:kaede)*, n=15; *MZsap130a;Tg(nkx2.5:kaede)* NV, n=14; *MZsap130a;Tg(nkx2.5:kaede)* SV, n=10. Scale bar 50μm

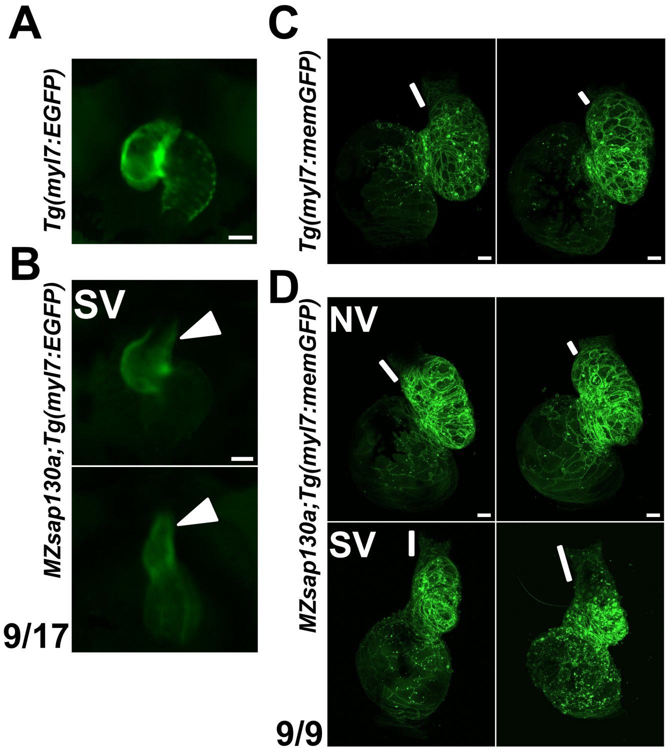


**FigS7:** ***MZsap130a*** **OFT images at 48-72hpf**

(**A, B**) *Tg(myl7:EGFP*) and *MZsap130a;Tg(myl7:EGFP)* embryos at 48hpf with longer OFTs. (**C, D**) *Tg(myl7:memGFP*) and *MZsap130a;Tg(myl7:memGFP)* embryos at 72hpf with longer OFTs. White triangles and lines highlight OFT structures. Scale bar = 50μm

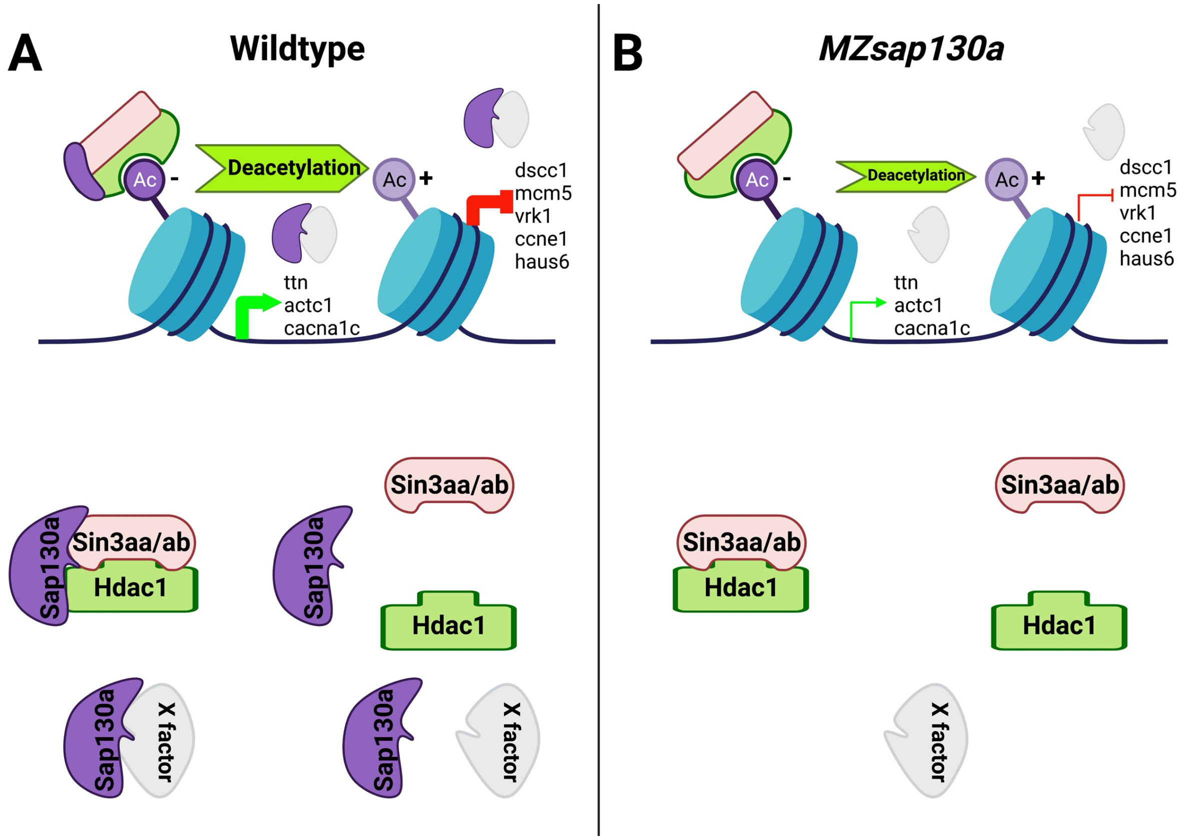


**FigS8: Proposed role for *sap130a* in *sin3aa*/*ab* and *hdac1* complex in zebrafish**
(**A**) Shows wildtype setting with Sap130a present interacting with the Sin3aa/ab Hdac1 complex or an unknown X factor. (**B**) Shows the possibilities when Sap130a is missing.
